# RF-Net 2: Fast Inference of Virus Reassortment and Hybridization Networks

**DOI:** 10.1101/2021.05.05.442676

**Authors:** Alexey Markin, Sanket Wagle, Tavis K. Anderson, Oliver Eulenstein

**Affiliations:** Virus and Prion Research Unit, National Animal Disease Center, USDA-ARS, USA; Department of Computer Science, Iowa State University, USA

## Abstract

**Motivation:** A phylogenetic network is a powerful model to represent entangled evolutionary histories with both divergent (speciation) and convergent (e.g., hybridization, reassortment, recombination) evolution. The standard approach to inference of hybridization networks is to (i) reconstruct rooted gene trees and (ii) leverage gene tree discordance for network inference. Recently, we introduced a method called *RF-Net* for accurate inference of virus reassortment and hybridization networks from input gene trees in the presence of errors commonly found in phylogenetic trees. While RF-Net demonstrated the ability to accurately infer networks with up to four reticulations from erroneous input gene trees, its application was limited by the number of reticulations it could handle in a reasonable amount of time. This limitation is particularly restrictive in the inference of the evolutionary history of segmented RNA viruses such as influenza A virus (IAV), where reassortment is one of the major mechanisms shaping the evolution of these pathogens.

**Results:** Here we expand the functionality of RF-Net that makes it significantly more applicable in practice. Crucially, we introduce a fast extension to RF-Net, called *Fast-RF-Net*, that can handle large numbers of reticulations without sacrificing accuracy. Additionally, we develop automatic stopping criteria to select the appropriate number of reticulations heuristically and implement a feature for RF-Net to output error-corrected input gene trees. We then conduct a comprehensive study of the original method and its novel extensions and confirm their efficacy in practice using extensive simulation and empirical influenza A virus evolutionary analyses.

**Availability:** *RF-Net 2* is available at https://github.com/flu-crew/rf-net-2.

## 1 Introduction

Phylogenetic analyses have made significant contributions towards our understanding of how genes, genomes, and species have evolved. The application of these techniques are far-reaching, affecting conservation biology, agriculture, drug development, epidemiology, and pandemic preparedness (Grenfell *et al*., 2004; Harris *et al*., 2013; Nik-Zainal *et al*., 2012; Jackson, 2004; Garten *et al*., 2009). However, when evolutionary processes such as hybridization, recombination, and horizontal gene transfer are involved, a non-treelike evolutionary history emerges. Omitting these processes may lead to incorrect inference (Posada and Crandall, 2002; Woolley *et al*., 2008), and can confound attempts to apply methods to solve real-world problems, e.g., reconstructing epidemic history and describing virus transmission dynamics (Leitner, 2019). Consequently, more complex evolutionary models in the form of phylogenetic networks have been developed that explicitly model reticulate evolutionary events (see reviews in Posada and Crandall, 2001; Huson and Bryant, 2006).

Phylogenetic networks model evolutionary history by adapting standard tree models (e.g., rooted binary trees) to also include *reticulation* events (Figure 1, left). Reticulation nodes can capture hybridization and similar convergent processes (Huson *et al*., 2010). Existing algorithms have significant promise and it is a rapidly growing field with some recent success in application to important biological questions. For example, Wen *et al*., 2016 developed new reticulation network methods that quantified incomplete lineage sorting and introgression in the malaria vector; and Willyard *et al*., 2009 applied network approaches to address questions of hybrid ancestry. Recently, Markin *et al*., 2019 introduced a method that adapted the standard hybridization network problem to address erroneous input trees using the classic Robinson-Foulds distance. This method had the utility of being able to objectively embed input gene trees into reticulation networks inexactly, but up to a measurable error, and was able to accurately infer phylogenetic networks while directly accounting for errors in the input gene trees. Unfortunately, while the Markin *et al*., 2019 method, RF-Net, can handle hundreds of taxa, it cannot address networks with large numbers of reticulations (e.g., greater than 10); a limitation common to most other network inference methods (Wen *et al*., 2016; Markin *et al*., 2019).

**Figure 1:**
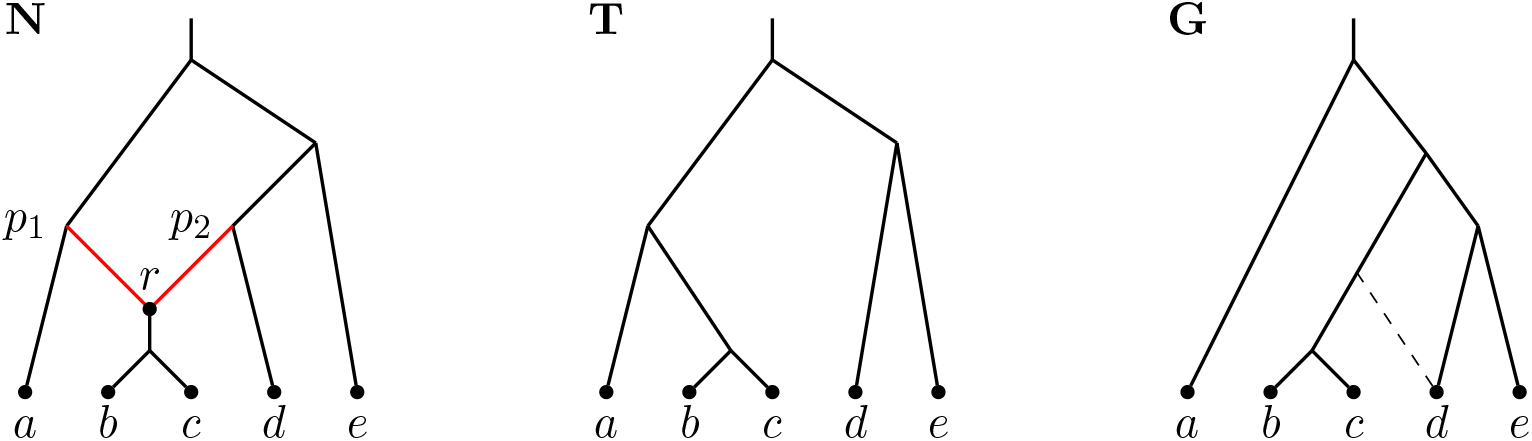
From left to right: Network *N*, tree *T* displayed in *N*, and tree *G. N* contains one reticulation vertex *r* with the entering reticulation edges shown in red. *T* is displayed in *N* by removing edge (*p*_2_, *r*). While *G* is not directly displayed in *N, N* displays a local modification of *G* indicated by the dashed edge.

The inability to model extensive reticulation in phylogenetic networks is particularly problematic in viruses. A reticulation process, frequently referred to as “reassortment”, has been reported to occur in at least 11 RNA virus families, and is a major driver in the evolution of pathogens such as influenza A virus (IAV), rotavirus A, and bluetongue virus (McDonald *et al*., 2016). In these viruses, reassortment occurs when two single-stranded segmented RNA viruses infect the same cell and exchange complete gene segments (McDonald *et al*., 2016). In an observational study that tracked gene combinations in H1 swine IAV, and consequently the minimum number of reassortments (i.e., unique gene pairings are derived from reassortment), Gao *et al*., 2017 detected more than 70 unique genome constellations. Similarly, Rajão *et al*., 2017 identified more than 44 unique genome constellations in swine H3 IAV. In avian IAV, reassortment is also ubiquitous (Lu *et al*., 2014).In certain locations, the ecology and diversity of the avian IAV appears to facilitate extensive reassortment and the emergence of novel viruses, e.g., Venkatesh *et al*., 2018 describe at least 24 HA/NA gene pairings across 12 HA subtypes. Though many of these reassorted viruses appear to be infrequently detected, when viral progeny contain genes derived from more than one parent, it may confer important fitness advantages (Scholtissek, 1990; Scholtissek *et al*., 1985; Anderson *et al*., 2020; Crisci *et al*., 2013). The most notable example how a reassorted virus can have far reaching impact is the 2009 H1N1 influenza pandemic virus. This lineage of IAV was the result of a reassortment event (or events) in swine that generated an IAV that contained gene segments with evolutionary origins from bird, human, and swine IAV (Smith *et al*., 2009). Consequently, given the importance of reassortment in virus biology, e.g., (Forster *et al*., 2020; Mengual-Chuliá *et al*., 2016), techniques that identify reassorted genomes, their evolutionary history, and allow for large numbers of reticulation vertices are critical.

### Related work

Phylogenetic networks have been broadly studied from many different perspectives – see Huson *et al*., 2010; Huson and Scornavacca, 2011 for a comprehensive review. A major focus in the literature is on the inference of hybridization networks (cf. a review in Elworth *et al*., 2018). The original approach to hybridization network inference was formulated by Baroni *et al*., 2005, which leveraged gene tree discordance. The original Baroni et al. work was followed by multiple algorithmic advancements (cf. Whidden *et al*., 2013; Albrecht, 2016; Iersel *et al*., 2014). Further, Meng and Kubatko, 2009; Yu *et al*., 2012 introduced the multi-species network coalescent (MSNC) model that combined the hybridization framework of Baroni et al. with the classic multi-species coalescent model (Kingman, 1982), which accounts for incomplete lineage sorting in the gene trees. This led to the development of multiple network inference approaches based on MSNC (cf. Solís-Lemus and Ané, 2016; Yu *et al*., 2012; Yu and Nakhleh, 2015). Additionally, Yu *et al*., 2013 proposed a parsimony method, MP-PhyloNet, for the inference of hybridization networks in the presence of incomplete lineage sorting.

Recently, RF-Net (Markin *et al*., 2019) extended the Baroni et al. hybridization framework to account for errors in the input gene trees (Steel and Rodrigo, 2008). Accounting for errors is crucial in many biological systems, where evolutionary inference is error-prone, e.g., (Hahn, 2007; Rasmussen and Kellis, 2011) and the classic incomplete lineage sorting model (the multi-species coalescent) is not entirely applicable, e.g., RNA viruses. To accommodate potential errors in gene trees, Markin *et al*., 2019 introduced the concept of *embeddings* and *embedding cost*. Given a gene tree *G* and a network *N*, an embedding of *G* in *N* is a tree *G*′ that is *displayed* in *N* which is most similar to *G* (i.e., *G*′ can be obtained from *N* by removing some reticulation edges). If *N* is the *true* phylogenetic network, then we can expect the embedding *G*′ to be the error-corrected version of *G*. The embedding cost is then the Robinson-Foulds distance between *G* and its embedding. RF-Net uses the sum of embedding costs over all gene trees as its objective function to perform local search on the space of phylogenetic networks. The local search is driven by the SubNet Prune and Regraft (SNPR) edit operation introduced by Bordewich *et al*., 2017. Note that SNPR was proven to ‘connect’ the space of phylogenetic networks (Janssen *et al*., 2018; Bordewich *et al*., 2017).

### Our contribution

In this work, we expand the functionality of RF-Net (Markin *et al*., 2019) to advance its practical applicability. We develop the *Fast-RF-Net* approach that is not bound to a limited number of reticulations which resolves a major drawback in the base RF-Net method. The core idea behind Fast-RF-Net is that, as RF-Net adds a new reticulation to a candidate phylogenetic network, different genes are likely to ‘favor’ one reticulation path to another based on the respective gene tree topology. Subsequently, as the search gradually progresses, we hypothesize that these preferences are unlikely to change, as each added reticulation improves the fitness of the network. This implies that we do not have to recompute gene tree embeddings ‘from scratch’ for every new candidate network, but can re-use the previous embeddings.

Second, we address a common problem in phylogenetic network inference and develop an approach that can determine the appropriate number of reticulations for a given dataset (Cai and Ané, 2020; Hejase *et al*., 2018). In Section 3.3 we propose an automated stopping criterion for RF-Net that does not require a user-specified number of reticulations. Additionally, RF-Net can plot the dynamic of the fitness function (embedding cost) as it changes with number of reticulations, which provides practitioners a visualization to aid in preventing over-fitting and selecting the best inferred network (Hejase *et al*., 2018).

We then conduct a comprehensive study of RF-Net and its novel components on (i) a small validation IAV dataset with clear patterns of reassortment (Section 4); (ii) simulated data (Section 5); and (iii) a large swine H3 IAV dataset with 429 whole genome strains and highly incongruent gene evolution (Section 6).

The simulation study demonstrated the ability of Fast-RF-Net to match the accuracy of RF-Net on datasets with up to 10 reticulations. Further, the simulations demonstrated that both RF-Net and Fast-RF-Net had a robust error-correction ability. The validation study showed that RF-Net was able to accurately recapitulate the evolutionary history of swine IAV, and the associated real-world empirical analyses revealed important evolutionary dynamics of the virus lineage including the identification of reassorted viruses that have pandemic potential.

## 2 Background

### 2.1 Preliminaries

#### Phylogenetic networks

A *(phylogenetic) network* is a directed acyclic graph (DAG) with a designated root vertex of in-degree zero and out-degree one and all other vertices are either of in-degree one and out-degree two (*tree vertices*), in-degree two and out-degree one (*reticulation vertices*), or in-degree one and out-degree zero (*leaves*). An example network with one reticulation vertex *r* is depicted in Figure 1 (right).

Let *N* be a network, then its vertices, edges, root, and leaves are denoted by *V* (*N*), *E*(*N*), *ρ*(*N*), and L(*N*), respectively. The edges in *E*(*N*) are distinguished by the edges that are entering (i) reticulation vertices (*reticulation edges*) and (ii) tree vertices or leaves (*tree edges*). A *tree-path* in *N* is a directed path that consists only of tree edges.

A vertex *v* ∈ *V* (*N*) is a *descendant* of *w* ∈ *V* (*N*) when there is a directed path from *w* to *v* (we consider each vertex to be a descendant of itself). A *(hardwired) cluster* of vertex *v, C*_*v*_, is the set of leaves that are descendants of *v*. Finally, 𝒞 (*N*) is the set of all clusters in *N*.

#### Phylogenetic trees

A *(phylogenetic) tree T* is a network with no reticulation vertices. Given a set *L* ⊆ L(*T*), *T* |_*L*_ denotes a restriction of *T* to the leaf-set *L*. That is, *T* |_*L*_ is obtained from the minimal subtree *T′* of *T*, which connects all leaves in *L* and the root, by suppressing all out-degree 1 nodes except for the root.

#### Displayed trees

Tree *T* is *displayed* in network *N* (with the same leaf set) if one can remove exactly one reticulation edge from each reticulation node, remove all potentially appearing non-labeled vertices with out-degree zero, and obtain a subdivision of *T*. Figure 1 demonstrates an example of tree *G* (left) displayed in network *N* (right).

The set of all trees displayed by *N* is denoted by 𝒫 _*N*_.

#### Tree-child networks

A network is called *tree-child* if each non-leaf vertex has at least one outgoing tree edge (i.e., a child that is a tree-vertex). Observe that each vertex in a tree-child network must have a tree-path going to some leaf.

#### Robinson-Foulds (RF) distance

The Robinson-Foulds distance between two trees is defined as the symmetric difference of their cluster representations (Robinson and Foulds, 1981). In particular,

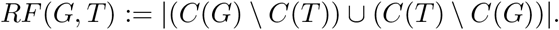

When one of the trees has an incomplete set of taxa, as is common in super-tree/network studies, we apply the standard *minus-method* approach (Cotton and Wilkinson, 2007). That is, if L(*G*) ⊂ L(*T*), we have

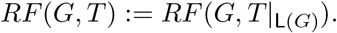

### 2.2 RF-networks

We now briefly re-iterate the definition of RF-Networks and the respective optimization problem from (Markin *et al*., 2019). We first introduce our core ‘fitness’ function for phylogenetic networks and then formulate the respective optimization problem.

#### Embeddings

To score the ‘fitness’ of network *N* against tree *G*, we need to consider the trees displayed in *N*. In particular, we define the *embedding cost* as follows (recall that 𝒫_*N*_ denotes the set of all trees displayed in *N*)

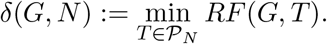

A tree *T* displayed in *N* with *RF* (*G, T*) = *δ*(*G, N*) is called an *embedding* of *G* in *N*. Note that the leaf-set of *G* should be a subset of the leaf-set of *N*.

As an example, consider Figure 1. *T* in that example is displayed in *N* and therefore *δ*(*T, N*) = 0. At the same time tree *G* is not displayed in the network, while a small modification of *G* indicated using the dashed edge is displayed; let us denote this modified tree as *G* ′. It is then not difficult to see that *δ*(*G, N*) = *RF* (*G, G*′) = 2.

Then, for a set of input trees 𝒢 and a network *N* (with L(*G*) ⊆ L(*N*) fo r all *G* ∈ 𝒢) the *total embedding cost* is the sum of individual embedding costs. That is, 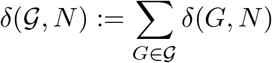.

Markin *et al*., 2019 demonstrated that computing *δ*(*G, N*) is NP-hard even when *N* is tree-child. This means that computing the embedding cost exactly when the number of reticulations is high (i.e., greater that 10 − 11) is computationally impractical for datasets with hundreds of taxa (see Section 3.2).

#### Optimization problem

We now formulate our core optimization problem that we approach with RF-Net.

**Problem 1**. RF-Network

*Input:* A set of input trees 𝒢 and the maximum number of reticulations *r*;

*Output:* Find network *N* with at most *r* reticulations minimizing the total embedding cost *δ*(𝒢, *N*). Note that *N* should contain all leaves (taxa) from the input trees

Note that if we set *r* to 0, this problem reduces to the well-studied Robinson-Foulds supertree problem (Bansal *et al*., 2010) which is known to be NP-hard (McMorris and Steel, 1994). Consequently, the RF-Network problem is also NP-hard.

## 3 Methods

We now describe *RF-Net* and introduce its novel extensions. We start with the base local search methodology; then we describe our key extension that enables RF-Net to handle a very large number of reticulations (*Fast-RF-Net*). Finally, we describe additional advancements that strengthen the applicability of RF-Net.

### 3.1 Method summary

Markin *et al*., 2019 proposed a ‘layered’ approach to RF-Network inference. In particular, we call a set of all networks with exactly *k* reticulations a *k-th layer* and it is denoted by 𝒩 _*k*_. Then the optimization search starts with the 0-th layer (i.e., tree layer) and then incrementally explores each higher layer until it reaches the user-specified upper bound on the number of reticulations, *r*. More precisely, the method first finds a locally optimal supertree *N* ^0^ for the input gene trees. Recall that when *N* is a tree, the embedding cost is the Robinson-Foulds distance. Consequently, we use the highly efficient RF Supertree method by Bansal *et al*., 2010 for this step.

Next, RF-Net repeats the following procedure until it reaches the layer with *r* reticulations (or the total embedding cost is reduced to 0). It starts with *k* = 0.

i. Add a reticulation to *N*^*k*^ in a way that minimizes the total embedding cost. Let *N*^*k*+1^ denote the resulting network. If there are multiple optimal ways to add a reticulation, RF-Net randomly chooses *p* optimal *N*^*k*+1^ networks and performs step (ii) for each of them sequentially. Here *p* is a user-specified parameter.
ii. Explore layer 𝒩_*k*+1_ using SNPR edit operations Bordewich *et al*., 2017, starting with an *N*^*k*+1^ network. At each local search iteration, if there are multiple optimal networks in the SNPR neighborhood, RF-Net proceeds by choosing one of them uniformly at random.

Note that RF-Net implements two major algorithmic advancements as described in Markin *et al*., 2019. These algorithms (1) make the computation of the total embedding cost for each network significantly faster, and (2) accelerate SNPR neighborhood traversals for each local search iteration. Finally, RF-Net can constraint the network search space to tree-child networks only. As in practice many reticulate histories can be expected to be tree-child (Markin *et al*., 2019; Cardona *et al*., 2008), this setting can noticeably reduce the search space and, hence, the runtime of RF-Net.

### 3.2 Fast-RF-Net: Scaling to highly entangled datasets

Despite the effective optimizations proposed in Markin *et al*., 2019, the computational hardness of calculating the RF embedding cost prevents RF-Net from inferring networks with more than 10-11 reticulations in a reasonable time (for larger datasets with 300-500 taxa). To overcome this in practice stringent limitation, we now describe a novel fast extension to RF-Net that can scale to an arbitrary number of reticulations.

Let *n*_1_, *n*_2_, …, *n*_*k*_ denote the reticulation vertices in network *N*^*k*^ in the *k*-th layer. For each input tree *G*, RF-Net computes an embedding in *N*^*k*^. That is, for each reticulation vertex *n*_*i*_, it chooses one of it’s parent-edges for the embedding. We can then represent an embedding for *G* as (*n*_1_*/p*_1_, *n*_2_*/p*_2_, …, *n*_*k*_*/p*_*k*_), where *p*_*i*_ ∈ {1, 2} represents whether the ‘first’ or the ‘second’ parent of *n*_*i*_ is chosen (or left/right).

Then during the traversal of the SNPR neighborhood of *N*^*k*^, to evaluate a network *N* ′, Fast-RF-Net uses the same (*n*_1_*/p*_1_, *n*_2_*/p*_2_, …, *n*_*k*_*/p*_*k*_) embedding for *G* instead of recomputing the optimal embedding from scratch. That way, the embedding cost for *N* ′ can be computed in *O*(| L(*N* ′)| · | 𝒢 |) time instead of *O*(2^*k*^| L(*N* ′)| · | 𝒢 |) time required by RF-Net.

Similarly, when adding a new reticulation vertex *n*_*k*+1_, Fast-RF-Net maintains the previous parent assignments (*n*_1_*/p*_1_, *n*_2_*/p*_2_, …, *n*_*k*_*/p*_*k*_) for *G* and then chooses the best parent assignment for *n*_*k*+1_ between *n*_*k*+1_*/*1 and *n*_*k*+1_*/*2 by computing the respective RF distances.

In summary, Fast-RF-Net eliminates the exponential factor 2^*k*^ from the RF-Net’s time complexity as a trade-off for potentially considering sub-optimal embeddings.

### 3.3 Determining the number of reticulations

It is generally hard (or even infeasible) to estimate the exact number of hybridization/reassortment events that took place in the past for a particular dataset. Therefore, to assist practitioners with model selection and determining the proper number of reticulations, we implemented an automated heuristic and a supplementary visual tool as described below.

#### 3.3.1 Analyzing the embedding cost dynamic

To aid practitioners, RF-Net can plot the dynamic of the total embedding costs; i.e., its decrease as the number of reticulations grows. Such a plot can then help identify the breakpoint, where RF-Net stops picking on the main reassortment/hybridization signal, but instead starts ‘over-fitting’ based on the topological error in gene trees (noise).

#### 3.3.2 Automated stopping criteria

Following the similar idea to Section 3.3.1, we propose a stopping criterion to determine the proper number of reticulations heuristically.

Let *C*_0_ denote the initial embedding cost (the total RF distance) for *N* ^0^. We then introduce a (percentage) parameter 0 ≤ *t <* 100 and require a difference between the embedding costs for two neighboring layers to be at least 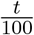 · *C*_0_.

More formally, let *F*_*k*_ denote the final network for a *k*-th layer. Then 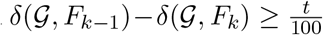 for all layers that RF-Net reports in the output. Once the improvement on the next layer is lower than the threshold, the search terminates. From now on we will refer to *t* as a *reticulation threshold*.

### 3.4 Error-correcting gene trees

Recall that computation of the embedding costs allows RF-Net to account for errors in the input gene trees. Consequently, in addition to a reassortment/hybridization network, RF-Net can produce the respective embeddings of the gene trees into the network. That is, RF-Net can output the error-corrected gene trees, which we empirically confirm in the simulation study (see Section 5).

When the embedding of an input tree into the computed network is not unique, RF-Net outputs a strict consensus of all embeddings for such input tree. On the other hand, Fast-RF-Net maintains a single embedding structure throughout the entire search and, therefore, outputs that particular embedding.

## 4 Validation of RF-Net with swine influenza A virus data

To evaluate the efficacy of RF-Net at inferring phylogenetic networks and accurately recreating reassortment events, we apply it to a small dataset of swine IAVs that include strains with evidence for reassortment based upon gene tree incongruence (Boni *et al*., 2010). We collated hemagglutinin (HA) and neuraminidase (NA) gene sequences for 22 swine IAV strains collected in the US as part of a national surveillance program (Anderson *et al*., 2013). These data included 21 strains from the H1N2 subtype within the delta-1a HA genetic clade: these HA genes were paired with a N2-1998 lineage or N2-2002 lineage NA gene. We included a single strain from the H1N1 subtype that had the same delta-1a HA genetic clade, but was paired with a Classical swine lineage N1 NA gene. From these data, we aligned each gene dataset with default settings in mafft v7.453 (Katoh *et al*., 2005), and inferred maximum likelihood gene trees for the separate alignments using FastTree v2.1.11 (Price*et al*., 2010). We then rooted the inferred trees with TreeTime v0.7.5 (Sagulenko *et al*., 2018).

The NA gene tree, Figure 2, demonstrates three different clades represented by the branch colors (red, green, and blue) reflecting the two subtypes and three evolutionary lineages. The HA gene tree reflects a single evolutionary lineage, but when the branches are colored by NA gene, distinct reassortment patterns are evident. First, while the H1N1 strain (blue) is nested within the HA gene tree, it is a distinct outgroup lineage in the NA tree. Second, in the NA gene tree, we note two large monophyletic clades reflecting the two distinct N2 lineages, N2-1998 vs N2-2002: in the HA tree, the N2-1998 green strains are nested within N2-2002 red strains. Taken together, this topological gene tree incongruence supports the proposition that at least two reassortment events have occured during the evolution of these swine IAV strains (Boni *et al*., 2010).

**Figure 2:**
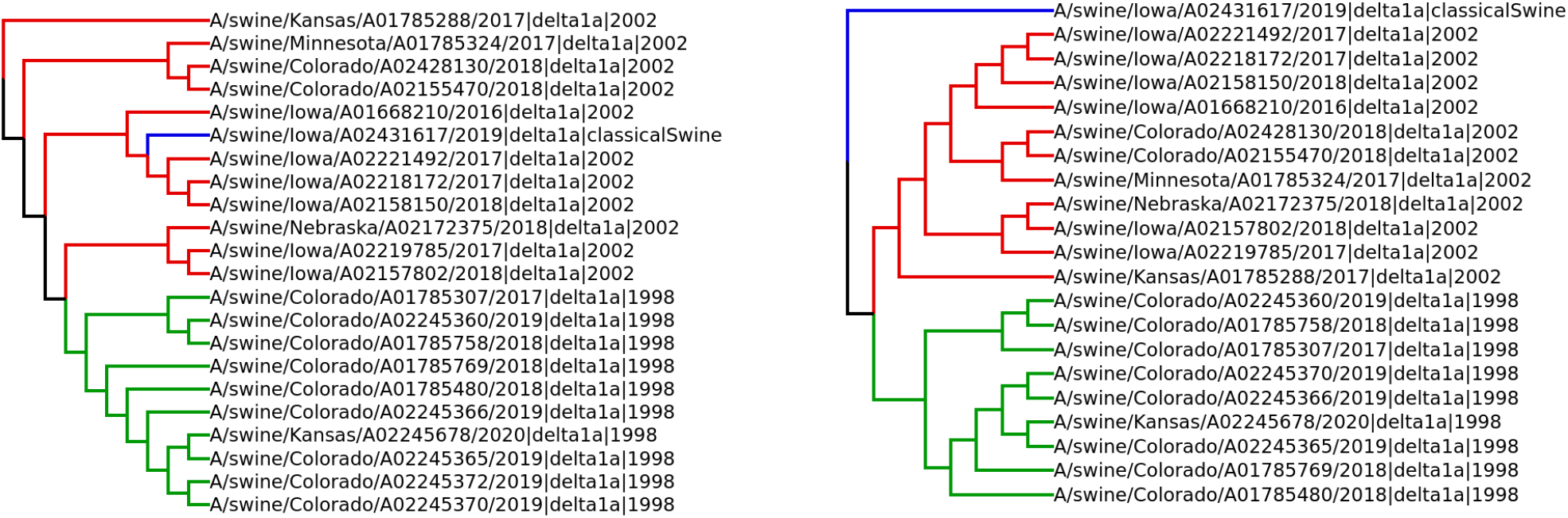
Evolutionary history of swine influenza A virus H1 delta-1a HA genes (left) with paired N1 or N2 genes (right). Influenza A virus strains are annotated by strain name, HA genetic clade, and NA genetic clade. The NA genetic clades are highlighted by blue, red, and green colors to represent N1-Classical swine, N2-2002, and N2-1998 clades, respectively

We subsequently conducted an RF-Net analysis to infer a network with two reassortments with the rooted HA and NA gene trees as input. Figure 3 represents the resulting inferred network. RF-Net correctly inferred the reassortment event at the H1N1 strain, allowing the network to reflect the position of that strain both in the HA and NA gene trees. RF-net was also able to infer the reassortment event demonstrated by the N2-1998 green strains, with a dashed line indicating the acquisition of this novel N2-1998 gene. Hence, we observe that RF-Net is able to recreate expected reassortment events with a high degree of resilience to small errors or inconsistencies in the input trees (e.g., derived from sequence error, alignment error, or lack of phylogenetic signal in highly similar IAV gene sequences).

**Figure 3:**
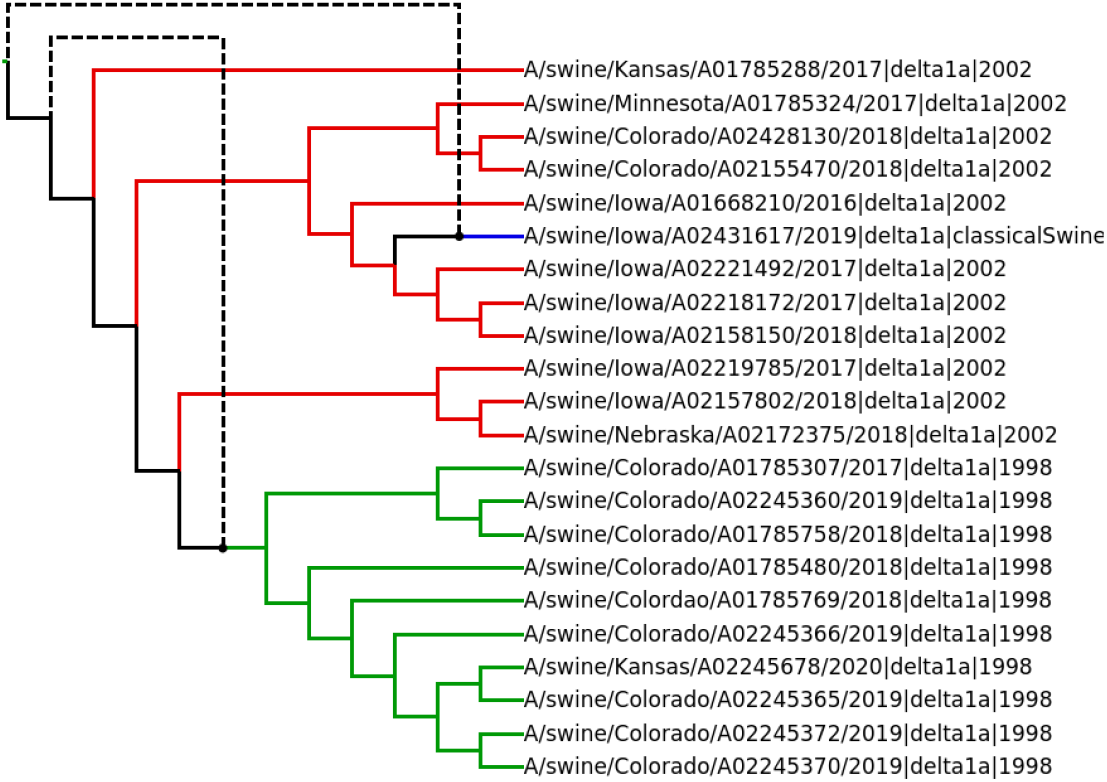
RF-Net network demonstrating two inferred reassortment events represented by dashed lines. RF-Net correctly inferred the novel HA and NA gene pairing of the H1N1 strain, and detected a reassortment within the H1 delta-1A lineage gene where reassortment resulted in a novel gene pairing of H1 delta-1a HA with a N2-1998 gene versus the dominant N2-2002 gene.

## 5 Simulation study

We evaluate RF-Net in a simulation study using the credible model of influenza A virus evolution from Müller *et al*., 2020. In this study, we deliberately use erroneous gene trees to evaluate RF-Net’s ability to perform error-correction. Within this simulation study we also compare the base RF-Net method with the Fast-RF-Net implementation from Section 3.2.

### 5.1 Simulation setup

Recently, Müller *et al*., 2020 introduced a statistical model of influenza A virus evolution by augmenting the classical coalescent model (Kingman, 1982) with reassortment. We apply their model, referred to as *coalescent with reassortment (CoalRe)*, to simulate reticulation networks and respective gene trees following the process described in Müller *et al*., 2020. We also simulated sequence alignments given the true gene trees, estimate (erroneous) gene trees from the alignments, and execute RF-Net on the estimated gene trees.

#### Simulating networks and gene trees

The CoalRe model uses the following parameters:

- Rate of reassortment per lineage per year. That is, in CoalRe, each lineage (going backward in time) has a fixed probability, *ρ*, of becoming a product of reassortment;
- Population size (a standard parameter in coalescent models).

We then use the following parameters in CoalRe simulations:

i. 50 and 75 for the number of taxa;
ii. a normal distribution with mean 0.05 and standard deviation 0.01 for the rate of reassortment;
iii. a normal distribution with mean 5 and standard deviation 0.1 for the population size;
iv. a uniform distribution over the [0, 10] interval as the sampling time (in years) for the taxa.
v. we set the number of genes to 8, matching the number of genes in influenza A virus. That is, each reassortment network is accompanied by 8 gene trees displayed in it.

Note that we choose the mean value of 5 for the population size, as it corresponds to the population size estimates for human IAVs by Müller *et al*., 2020. The chosen reassortment rate allowed us to obtain networks with approximately 4 reassortments for 50 taxa and 6 reassortments for 75 taxa. The maximum number of reassortments in the simulated networks was 10.

#### Sequence alignments and estimated gene trees

Following Müller *et al*., 2020, we generated nucleotide sequence alignments of length 1000 with a 0.005 substitution rate per site per year for each gene. The chosen substitution rate aligns with the estimates in Zeller *et al*., 2020 for two major gene segments in swine IAVs. The substitutions are driven by a Jukes-Cantor substitution model (Jukes *et al*., 1969).

With the simulated sequence alignments, we inferred gene trees using IQ-Tree v1.6.12 (Nguyen *et al*., 2015) implementing a Jukes-Cantor substitution model. Each inferred gene tree was rooted using TreeTime v0.7.5 (Sagulenko *et al*., 2018). Importantly, each process in this analysis – though standard in phylogenetics (Dereeper *et al*., 2008; Kapli *et al*., 2020) – unintentionally introduces a substantial number of errors to the estimated gene trees.

#### Executing RF-Net

We generated 10 instances with 50 taxa and 10 more instances with 75 taxa (each instance represents a network with 8 respective true gene trees and 8 estimated gene trees). We then executed 6 variations of RF-Net on each of the instances using the estimated (erroneous) gene trees as input. Specifically, we conducted runs with:

(i) RF-Net with the true number of reassortments taken from the true network;
(ii) RF-Net with an automated stopping criterion – 5% reticulation threshold (see Section 3.3.2);
(iii) RF-Net with 3% reticulation threshold;
(iv-vi) Fast-RF-Net with same 3 stopping criteria.

For convenience, we denote the above 6 variations of the method as *RF-Net, t5, t3, Fast-RF-Net, Fast-t5, Fast-t3* respectively. Here t5 and t3 refer to the employed 5% and 3% reticulation thresholds, respectively.

The experiments were conducted on a standard Windows laptop with an Intel 2.5GHz CPU.

### 5.2 Simulation results

We evaluate RF-Net using two metrics. First, we measured the similarity of an inferred network to the true simulated network. Second, we measured how well RF-Net performs error-correction of the gene trees.

#### Network reconstruction accuracy

We evaluate the similarity between the reconstructed and true simulated networks using the generalized Robinson-Foulds (gRF) measurement (Cardona *et al*., 2008). In particular, we compute the similarity between two networks by calculating the number of *hardwired* clusters they have in common and then normalizing that value by the total number of hardwired clusters in the networks. The similarity value of 1 then represents the maximum similarity. Figure 4 shows the reconstruction accuracy of RF-Net (similarity to the true network) summarized over 10 replicates for 50 and 75 taxa. Notably, the figure shows that while, on average, Fast-RF-Net is consistently less accurate than RF-Net for all three stopping criteria, the difference in accuracy is generally quite small. Moreover, in several instances Fast-RF-Net proved to be more accurate than RF-Net.

**Figure 4:**
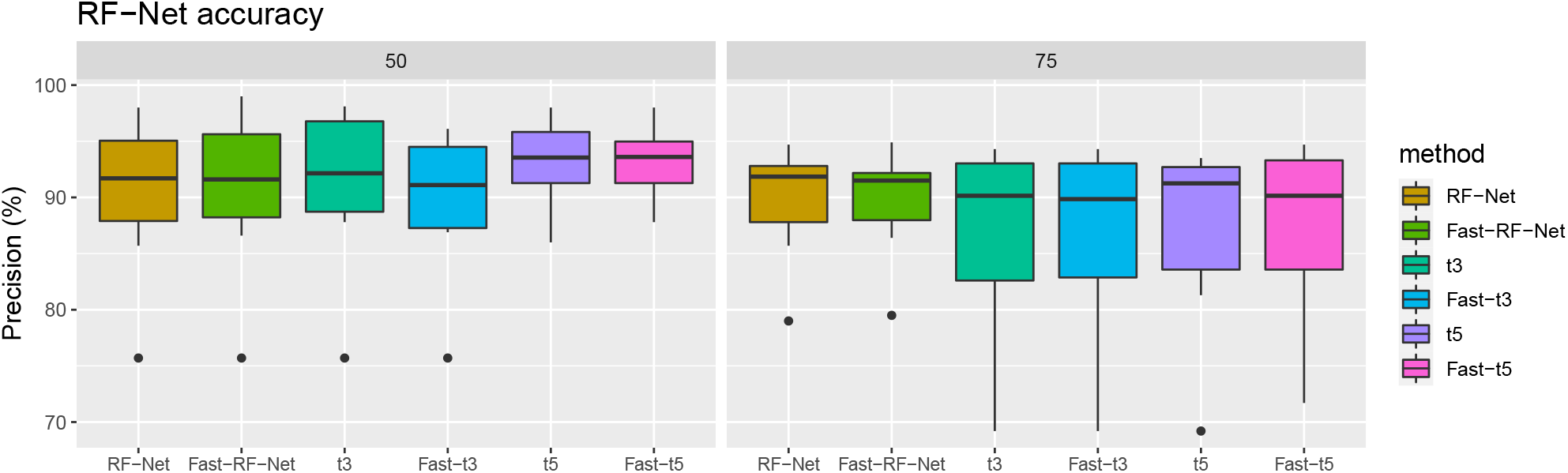
The precision of RF-Net variants, in terms of the similarity of the reconstructed networks with the respective true simulated networks, for 50 taxa (left) and 75 taxa (right).

Further, when focusing on 50 taxa instances, RF-Net with the automatic reticulation criteria (t3 and t5) was more accurate than when the true number of reassortments was given. This can be explained by the fact that CoalRe simulations often generate ‘parent-child’ reassortments on networks. In such a scenario, a reassortment vertex has two parents *p*_1_ and *p*_2_, where *p*_2_ is also a child of *p*_1_ (see Figure 6 for an illustration). Such reticulations cannot be detected from gene tree incongruence. Therefore, specifying the true number of reassortments will lead RF-Net to over-fitting. Hence, RF-Net with the automatic reticulation criteria demonstrated better accuracy.

We did not observe the same pattern for 75 taxa. In these simulations, specifying the true number of reassortments in RF-Net showed better accuracy on average. This is likely because parent-child reassortments are less frequent when the number of co-evolving strains is larger.

#### Evaluating error-correction

We evaluated the embeddings (see Section 3.4) of the input trees into an inferred network. Our main goal was to see whether the RF-Net embeddings were closer to the true simulated gene trees than the erroneous estimated gene trees. That is, we wanted to verify that RF-Net performs error-correction on the input trees.

The standard approach to measure error in trees is to count the number of incorrect clusters (false positives) and the number of missing clusters (false negatives). In our case the true simulated gene trees are always fully resolved (binary), whereas the embeddings can be not fully resolved. In these circumstances the number of false negatives is always at least as large as false positives. Therefore, we only considered the false negatives (as the worst case).

Figure 5 summarizes the performance of RF-Net in terms of error-correction, where errors are measured as the number of missing clusters. First of all, we observed that all variants of RF-Net effectively performed error-correction by significantly reducing the number of errors in the estimated gene trees. In fact, all variants reduced the number of errors in the estimated gene trees by more than 50% on average for both 50 and 75 taxon instances. For example, RF-Net with a reticulation threshold of 5% (t5) on average eliminated 64.0% of errors for 50 taxon instances and 51.2% of errors for 75 taxon instances. Finally, similarly to Figure 4, here we again observed only a small difference between RF-Net and Fast-RF-Net in terms of error-correction.

**Figure 5:**
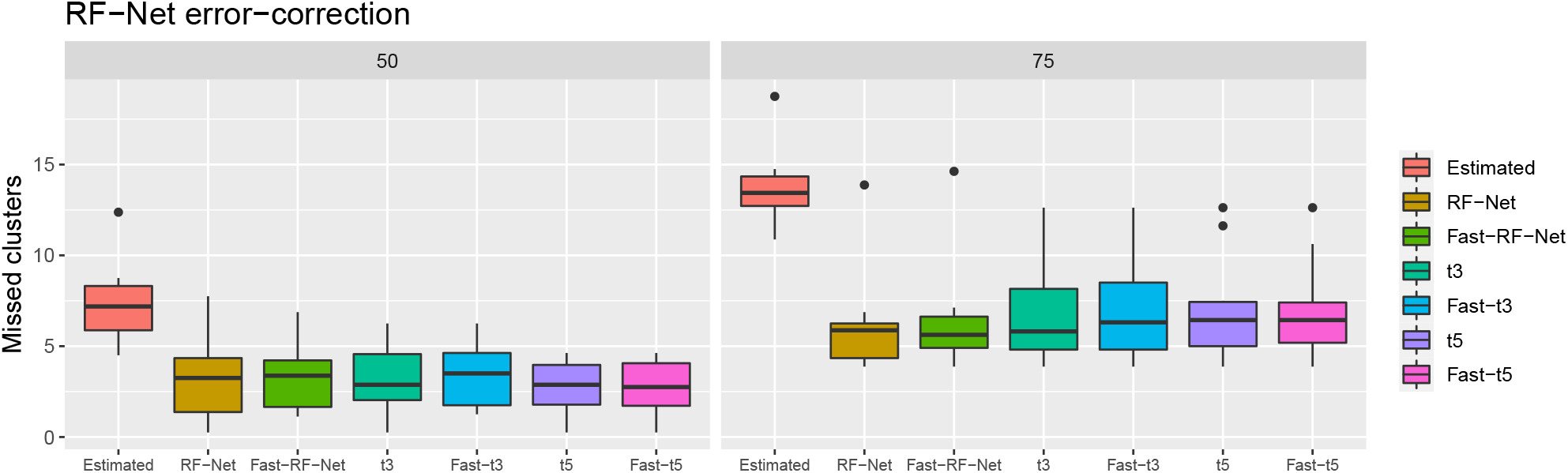
Number of missed clusters (false negatives) in the estimated gene trees and in the embeddings of the gene trees by tested variant of RF-Net.

**Figure 6:**
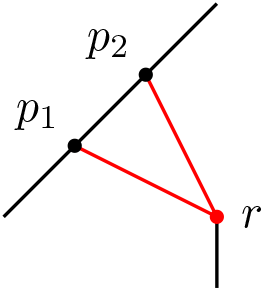
Example of a parent-child reticulation *r* with parents *p*_1_ and *p*_2_.

## 6 Empirical study

We demonstrate the utility of RF-Net, and Fast-RF-Net in particular, by applying it to an extensive swine IAV dataset and showcase the ability of RF-Net to identify novel reassorted viruses.

### Context and influenza A virus global relevance

The observed success of many pathogens likely depends on the coordinated function of combinations of genes. There is evidence that mutations in one gene that normally would result in a ‘dead’ pathogen that arise under drug treatment are alleviated by secondary compensatory mutations in other genes (Lozovsky *et al*., 2009; Weinreich *et al*., 2006; Wang *et al*., 2002). In IAV this is exemplified by the interplay between the two surface proteins – the hemagglutinin (HA) and neuramindase (NA). These genes function in a coordinated effort to replicate within and transmit between hosts: balance in NA and HA cellular interactions are necessary for IAV infection to result in successful host-to-host transmission (Mitnaul *et al*., 2000; Neverov *et al*., 2014; Das *et al*., 2011). Though there is little experimental work in swine IAV (but see examples in human IAV White and Lowen, 2018; White *et al*., 2019 and review by Lowen, 2017), there is observational evidence that certain gene combinations provide fitness advantages and disadvantages. This is evident from the presence of extensive reassortment of IAV in swine (Gao *et al*., 2017; Rajão *et al*., 2017; Diaz *et al*., 2017; Henritzi *et al*., 2020) with more than 100 distinct genome constellations, but only a small fraction of these constellations are frequently detected, suggesting that the majority of these reassorted viruses may have little epidemiological relevance (Powell *et al*., 2020).

Despite this observation, understanding the dynamics of reassortment and whether there are linkages between different swine IAV genes can inform and help minimize the risk of swine-origin IAV being transmitted to humans. This is because all swine IAVs circulating in the US contain genes derived from reassortment between swine-, human-, and avian-origin viruses (Gao *et al*., 2017; Rajão *et al*., 2017; Anderson *et al*., 2020). A consequence is that the evolution of human-origin IAV in the swine host may result in swine viruses to which the human population may have little to no immunity. Further, the emergence of novel gene combinations may spur rapid evolution in the surface proteins of IAV resulting in additional drift (Zeller *et al*., 2020). This is concerning as following the spread of the swine-origin 2009 H1N1 pandemic virus in humans, there has been the regular annual introduction of this human seasonal H1N1 virus into pigs, resulting in a decade of reassortment between human and in endemic swine IAV (Nelson *et al*., 2015). The direct consequence of the constant process of reassortment is that many swine IAV viruses may contain “human-adapted” genes that could have pandemic potential (Anderson *et al*., 2020).

In this empirical example, we focus on an H3.2010.1 swine IAV lineage derived from a human-seasonal H3 introduction to swine (Anderson *et al*., 2020; Rajão *et al*., 2015). Generally, human IAV infection in swine results in low replication and rare pig-to-pig transmission, but some human-origin IAV become endemic, and this is typically associated with genetic differences from the precursor strain (Lewis *et al*., 2016; Rajão *et al*., 2018; Powell *et al*., 2020), or reassortment with endemic host-adapted viruses. The H3.2010.1 lineage follows this paradigm, and assessing reassortment in this lineage allows us to generate testable hypotheses on how expansion in the genomic diversity of IAV in swine populations may facilitate IAV capable of spillover from swine to humans.

### Data collection

To study the evolution of the H3.2010.1 virus lineage, we downloaded all swine H3N2 complete genomes (n=1563) from the Influenza Research Database (Zhang *et al*., 2017). We then concatenated the 8 genes for each strain and aligned the genomes using mafft v7.453 (Katoh *et al*., 2005) to identify and remove redundant strains with identical genomes. The genomes were then separated into the 8 constituent genes, and the genetic clade and evolutionary lineage was inferred using the octoFLU classification pipeline (Chang *et al*., 2019). The genomes that contained a H3.2010.1 HA gene were used for our study, and strains were annotated by NA and evolutionary lineage for the 6 internal genes. As our goal was to describe reassortment in circulating swine IAV, strains that reflected single outbreak events collected as part of active surveillance at agricultural fairs (Bowman *et al*., 2017; Nelson *et al*., 2016) were removed from subsequent analyses resulting in a final dataset of 429 H3 whole genomes. A single outgroup was included, A/swine/Guatemala/CIP049-IP040078/2010, a dead-end human-to-swine H3 spillover detected in 2010. To generate the required input trees for RF-Net, the 8 genes were aligned, and maximum likelihood phylogenetic trees were inferred following automatic model selection using IQ-TREE (Nguyen *et al*., 2015). Rooting was validated using Tree-Time (Sagulenko *et al*., 2018): these data revealed inconsistent rooting in the NS gene tree. RF-Net relies upon tree topology to accurately infer reassortment, consequently, the NS gene was excluded from inference. The evolutionary lineage of the NS gene was maintained through manual annotation of strains.

### Experimental setup

This study was conducted on the USDA-ARS SCINet Ceres high-performance computing cluster https://scinet.usda.gov. We executed Fast-RF-Net on 7 reconstructed gene tree topologies in the tree-child mode as reassortment networks for IAV surveillance data can be expected to be tree-child (Markin *et al*., 2019). Consequently, we obtained reassortment network estimates with up to 20 reticulations in under 24 hours.

### Results and Discussion

The expanded functionality of RF-Net enabled us to infer the evolutionary history of a virus that is shaped by clonal and non-clonal processes. Given the frequency of reassortment in IAV, methods that do not consider reticulation processes may result in error if there is a reliance on single-gene inference. In analyzing our H3 swine IAV data, we were able to track the evolution of a major H3 lineage in swine as it reassorted multiple times; with reassortment coinciding with the genetic lineage becoming a major component of swine IAV diversity in the US (Zeller *et al*., 2018).

Our analysis recapitulates the three major reassortment events (Rajão *et al*., 2015; Powell *et al*., 2020). Specifically, the initial case (A/swine/Missouri/A01476459/2012) contained a human seasonal H3 hemagglutinin (HA), N2 neuraminidase (NA), and internal genes derived from the 2009 pandemic H1N1 (H1N1pdm09). The second reassortment event resulted in viruses where the human N2-NA was replaced by a classical swine N1-NA. The third major reassortment event involved the exchange of the N1-NA to an N2-NA derived from endemic swine 2002 N2 genes, the inclusion of the Matrix (M) gene from H1N1pdm09, with the remaining internal genes associated with the endemic swine triple reassortant internal gene (TRIG) constellation (Powell *et al*., 2020). Given the known minimum number of reassortment events in this IAV lineage, our method adequately recreates its evolutionary history. The phylogenetic networks computed by RF-Net, with over 400 strains, are difficult to visualize, but are provided at https://github.com/flu-crew/rf-net-2 and may be interactively explored with the browser-based network viewer IcyTree (Vaughan, 2017).

A significant feature of the expanded RF-Net tool is the exploration of networks with different numbers of reticulations *r* and the inclusion of an automatic stopping criterion when the improvement in the embedding cost is negligible (*t*). In our dataset, we explored networks with *r* ranging from 0 to 20 and determined whether biologically plausible reassorted strains were detected (Figure 7). In doing so, we noted additional reassortment events: these were also supported using single-gene phylogenetic methods (i.e., topological incongruence (Boni *et al*., 2010)), with the most interesting occurring in recent swine strains (e.g., A/swine/Illinois/A02218757/2017 and A/swine/Pennsylvania/ A02218184/2017). These reassorted viruses have been associated with recent spillovers to the human population (see (Bowman *et al*., 2017; Anderson *et al*., 2020; Duwell *et al*., 2018)). We also noted an almost linear decrease in the embedding cost as we increased the number of reticulations present in the network. This suggests that the empirical data best fit in networks with at least 20 reticulations (Figure 7). This is supported by empirical single-gene phylogenetic studies that demonstrated considerable amounts of reassortment; Rajão *et al*., 2017 detected at least 40 reassorted genomes in a different swine H3 IAV lineage. Our data also reveal what appear to be a number of intralineage reassortment events, where highly similar IAV coinfect cells and exchange genes, resulting in novel progeny. The intralineage reassortment process would be difficult to detect with standard methods such as tanglegrams that track topological incongruence (Venkatesh *et al*., 2018; Zeller *et al*., 2020) as the evolutionary gene lineages would appear to be the same despite being acquired from different parents. The consequence of such high levels of intralineage reassortment is unknown, but we hypothesize that this process contributes to the expansion in the genetic diversity of IAV in swine in the US, within genes and genotype constellations (Neverov *et al*., 2015; Zeller *et al*., 2020).

**Figure 7:**
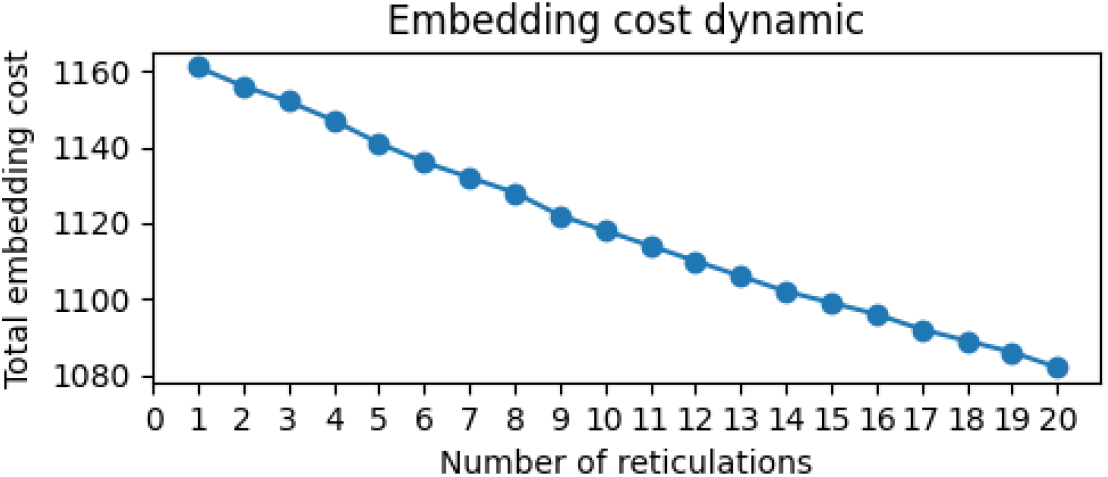
Dynamic decrease in embedding cost associated with number of reticulations in H3.2010.1 reticulation network.

## 7 Conclusion

We extended RF-Net with a suite of methods to advance its applicability in practice. Notably, the simulation study validated our hypothesis, implemented in Fast-RF-Net, that the reticulation paths in the embeddings of gene trees do not change significantly as the search progresses, which allowed us to scale RF-Net to large numbers of reticulations. Additionally, the simulations showcased the applicability of our proposed automated stopping criterion for the inference of the proper number of reticulate events. With the additional feature for the output of gene tree embeddings, we evaluated and confirmed the ability of RF-Net to successfully error-correct input trees. Finally, using Fast-RF-Net we were able to analyze the evolution of a major H3 swine IAV lineage with available whole genome data and observe many patterns of both inter- and intra-lineage reassortments. Importantly, the reassortment process has two biological outcomes: first, it increases the genetic and antigenic diversity of IAV within the swine ‘mixing vessel’ host (Scholtissek, 1990); and second, it increases the likelihood that a virus with zoonotic potential may emerge via a ‘sampling effect’, e.g., Loreau, 1998. Though our current analyses were post-hoc, RF-Net could be developed to run as an online algorithm, whereby surveillance data are rapidly screened for reassorted viruses, and novel detections could be screened in laboratory assays to preemptively determine human spillover potential.

## References

Albrecht, B. (2016). Computing hybridization networks using agreement forests. Ph.D. thesis, Ludwig-Maximilians-Universität München.

Anderson, T. K., Nelson, M. I., Kitikoon, P., Swenson, S. L., Korslund, J. A., and Vincent, A. L. (2013). Population dynamics of cocirculating swine influenza A viruses in the United States from 2009 to 2012. Influenza and Other Respiratory Viruses, 7, 42–51.

Anderson, T. K., Chang, J., Arendsee, Z. W., Venkatesh, D., Souza, C. K., Kimble, J. B., Lewis, N. S., Davis, C. T., and Vincent, A. L. (2020). Swine influenza a viruses and the tangled relationship with humans. Cold Spring Harbor perspectives in medicine, page a038737.

Bansal, M. S., Burleigh, J. G., Eulenstein, O., and Fernández-Baca, D. (2010). Robinson-Foulds supertrees. Algorithms Mol Biol, 5, 18.

Baroni, M., Semple, C., and Steel, M. (2005). A framework for representing reticulate evolution. Annals of Combinatorics, 8(4), 391–408.

Boni, M. F., de Jong, M. D., van Doorn, H. R., and Holmes, E. C. (2010). Guidelines for Identifying Homologous Recombination Events in Influenza A Virus. PLOS ONE, 5(5), 1–11.

Bordewich, M., Linz, S., and Semple, C. (2017). Lost in space? generalising subtree prune and regraft to spaces of phylogenetic networks. Journal of theoretical biology, 423, 1–12.

Bowman, A. S., Walia, R. R., Nolting, J. M., Vincent, A. L., Killian, M. L., Zentkovich, M. M., Lorbach, J. N., Lauterbach, S. E., Anderson, T. K., Davis, C. T., et al. (2017). Influenza A (H3N2) virus in swine at agricultural fairs and transmission to humans, Michigan and Ohio, USA, 2016. Emerging infectious diseases, 23(9), 1551.

Cai, R. and Ané, C. (2020). Assessing the fit of the multi-species network coalescent to multi-locus data. Bioinformatics.

Cardona, G., Llabrés, M., Rossellú, F., and Valiente, G. (2008). Metrics for phylogenetic networks i: Generalizations of the robinson-foulds metric. IEEE/ACM Transactions on Computational Biology and Bioinformatics, 6(1), 46–61.

Chang, J., Anderson, T. K., Zeller, M. A., Gauger, P. C., and Vincent, A. L. (2019). octoflu: automated classification for the evolutionary origin of influenza a virus gene sequences detected in us swine. Microbiology resource announcements, 8(32).

Cotton, J. A. and Wilkinson, M. (2007). Majority-rule supertrees. Syst Biol, 56(3), 445–452.

Crisci, E., Mussá, T., Fraile, L., and Montoya, M. (2013). Influenza virus in pigs. Molecular immunology, 55(3-4), 200–211.

Das, S. R., Hensley, S. E., David, A., Schmidt, L., Gibbs, J. S., Puigbú, P., Ince, W. L., Bennink, J. R., and Yewdell, J. W. (2011). Fitness costs limit influenza a virus hemagglutinin glycosylation as an immune evasion strategy. Proceedings of the National Academy of Sciences, 108(51), E1417–E1422.

Dereeper, A., Guignon, V., Blanc, G., Audic, S., Buffet, S., Chevenet, F., Dufayard, J.-F., Guindon, S., Lefort, V., Lescot, M., et al. (2008). Phylogeny. fr: robust phylogenetic analysis for the non-specialist. Nucleic acids research, 36(Suppl 2), W465–W469.

Diaz, A., Marthaler, D., Corzo, C., Muñoz-Zanzi, C., Sreevatsan, S., Culhane, M., and Torremorell, M. (2017). Multiple genome constellations of similar and distinct influenza a viruses co-circulate in pigs during epidemic events. Scientific reports, 7(1), 1–14.

Duwell, M. M., Blythe, D., Radebaugh, M. W., Kough, E. M., Bachaus, B., Crum, D. A., Perkins Jr, K. A., Blanton, L., Davis, C. T., Jang, Y., et al. (2018). Influenza a (h3n2) variant virus outbreak at three fairs—maryland, 2017. Morbidity and Mortality Weekly Report, 67(42), 1169.

Elworth, R. L., Ogilvie, H. A., Zhu, J., and Nakhleh, L. (2018). Advances in computational methods for phylogenetic networks in the presence of hybridization. arXiv preprint arXiv:1808.08662.

Forster, P., Forster, L., Renfrew, C., and Forster, M. (2020). Phylogenetic network analysis of sars-cov-2 genomes. Proceedings of the National Academy of Sciences, 117(17), 9241–9243.

Gao, S., Anderson, T. K., Walia, R. R., Dorman, K. S., Janas-Martindale, A., and Vincent, A. L. (2017). The genomic evolution of H1 influenza A viruses from swine detected in the united states between 2009 and 2016. Journal of General Virology, 98(8), 2001–2010.

Garten, R. J., Davis, C. T., Russell, C. A., Shu, B., Lindstrom, S., Balish, A., Sessions, W. M., Xu, X., Skepner, E., Deyde, V., et al. (2009). Antigenic and genetic characteristics of swine-origin 2009 A (H1N1) influenza viruses circulating in humans. Science, 325(5937), 197–201.

Grenfell, B. T., Pybus, O. G., Gog, J. R., Wood, J. L., Daly, J. M., Mumford, J. A., and Holmes, E. C. (2004). Unifying the epidemiological and evolutionary dynamics of pathogens. Science, 303(5656), 327–332.

Hahn, M. W. (2007). Bias in phylogenetic tree reconciliation methods: implications for vertebrate genome evolution. Genome biology, 8(7), 1–9.

Harris, S. R., Cartwright, E. J., Torök, M. E., Holden, M. T., Brown, N. M., Ogilvy-Stuart, A. L., Ellington, M. J., Quail, M. A., Bentley, S. D., Parkhill, J., and Peacock, S. J. (2013). Whole-genome sequencing for analysis of an outbreak of meticillin-resistant staphylococcus aureus: a descriptive study. Lancet Infect Dis, 13(2), 130–6.

Hejase, H. A., VandePol, N., Bonito, G. M., and Liu, K. J. (2018). Fastnet: fast and accurate statistical inference of phylogenetic networks using large-scale genomic sequence data. In RECOMB International conference on Comparative Genomics, pages 242–259. Springer.

Henritzi, D., Petric, P. P., Lewis, N. S., Graaf, A., Pessia, A., Starick, E., Breithaupt, A., Strebelow, G., Luttermann, C., Parker, L. M. K., et al. (2020). Surveillance of european domestic pig populations identifies an emerging reservoir of potentially zoonotic swine influenza a viruses. Cell Host & Microbe, 28(4), 614–627.

Huson, D. H. and Bryant, D. (2006). Application of phylogenetic networks in evolutionary studies. Molecular biology and evolution, 23(2), 254–267.

Huson, D. H. and Scornavacca, C. (2011). A survey of combinatorial methods for phylogenetic networks. Genome biology and evolution, 3, 23–35.

Huson, D. H., Rupp, R., and Scornavacca, C. (2010). Phylogenetic networks: concepts, algorithms and applications. Cambridge University Press.

Iersel, L. v., Kelk, S., Lekić, N., and Scornavacca, C. (2014). A practical approximation algorithm for solving massive instances of hybridization number for binary and nonbinary trees. BMC Bioinformatics, 15(1), 127.

Jackson, A. P. (2004). A reconciliation analysis of host switching in plant-fungal symbioses. Evolution, 58(9), 1909–23.

Janssen, R., Jones, M., Erdos, P. L., van Iersel, L., and Scornavacca, C. (2018). Exploring the tiers of rooted phylogenetic network space using tail moves. Bulletin of Mathematical Biology, 80(8), 2177–2208.

Jukes, T. H., Cantor, C. R., et al. (1969). Evolution of protein molecules. Mammalian protein metabolism, 3, 21–132.

Kapli, P., Yang, Z., and Telford, M. J. (2020). Phylogenetic tree building in the genomic age. Nature Reviews Genetics, 21(7), 428–444.

Katoh, K., Kuma, K.-i., Toh, H., and Miyata, T. (2005). Mafft version 5: improvement in accuracy of multiple sequence alignment. Nucleic acids research, 33(2), 511–518.

Kingman, J. F. C. (1982). The coalescent. Stochastic processes and their applications, 13(3), 235–248.

Leitner, T. (2019). Phylogenetics in hiv transmission: taking within-host diversity into account. Current Opinion in HIV and AIDS, 14(3), 181–187.

Lewis, N. S., Russell, C. A., Langat, P., Anderson, T. K., Berger, K., Bielejec, F., Burke, D. F., Dudas, G., Fonville, J. M., Fouchier, R. A., et al. (2016). The global antigenic diversity of swine influenza A viruses. Elife, 5, e12217.

Loreau, M. (1998). Separating sampling and other effects in biodiversity experiments. Oikos, pages 600–602.

Lowen, A. C. (2017). Constraints, drivers, and implications of influenza a virus reassortment. Annual review of virology, 4, 105–121.

Lozovsky, E. R., Chookajorn, T., Brown, K. M., Imwong, M., Shaw, P. J., Kamchonwongpaisan, S., Neafsey, D. E., Weinreich, D. M., and Hartl, D. L. (2009). Stepwise acquisition of pyrimethamine resistance in the malaria parasite. Proceedings of the National Academy of Sciences, 106(29), 12025–12030.

Lu, L., Lycett, S. J., and Brown, A. J. L. (2014). Reassortment patterns of avian influenza virus internal segments among different subtypes. BMC evolutionary biology, 14(1), 1–15.

Markin, A., Anderson, T. K., Vadali, V. S. K. T., and Eulenstein, O. (2019). Robinson-foulds reticulation networks. In Proceedings of the 10th ACM International Conference on Bioinformatics, Computational Biology and Health Informatics, pages 77–86. ACM.

McDonald, S. M., Nelson, M. I., Turner, P. E., and Patton, J. T. (2016). Reassortment in segmented rna viruses: mechanisms and outcomes. Nature Reviews Microbiology, 14(7), 448.

McMorris, F. and Steel, M. A. (1994). The complexity of the median procedure for binary trees. In New Approaches in Classification and Data Analysis, pages 136–140. Springer.

Meng, C. and Kubatko, L. S. (2009). Detecting hybrid speciation in the presence of incomplete lineage sorting using gene tree incongruence: a model. Theoretical population biology, 75(1), 35–45.

Mengual-Chuliá, B., Bedhomme, S., Lafforgue, G., Elena, S. F., and Bravo, I. G. (2016). Assessing parallel gene histories in viral genomes. BMC evolutionary biology, 16(1), 1–15.

Mitnaul, L. J., Matrosovich, M. N., Castrucci, M. R., Tuzikov, A. B., Bovin, N. V., Kobasa, D., and Kawaoka, Y. (2000). Balanced hemagglutinin and neuraminidase activities are critical for efficient replication of influenza a virus. Journal of virology, 74(13), 6015– 6020.

Müller, N. F., Stolz, U., Dudas, G., Stadler, T., and Vaughan, T. G. (2020). Bayesian inference of reassortment networks reveals fitness benefits of reassortment in human influenza viruses. Proceedings of the National Academy of Sciences, 117(29), 17104–17111.

Nelson, M. I., Stratton, J., Killian, M. L., Janas-Martindale, A., and Vincent, A. L. (2015). Continual reintroduction of human pandemic h1n1 influenza a viruses into swine in the united states, 2009 to 2014. Journal of virology, 89(12), 6218–6226.

Nelson, M. I., Wentworth, D. E., Das, S. R., Sreevatsan, S., Killian, M. L., Nolting, J. M., Slemons, R. D., and Bowman, A. S. (2016). Evolutionary dynamics of influenza a viruses in us exhibition swine. The Journal of infectious diseases, 213(2), 173–182.

Neverov, A. D., Lezhnina, K. V., Kondrashov, A. S., and Bazykin, G. A. (2014). Intrasubtype reassortments cause adaptive amino acid replacements in h3n2 influenza genes. PLoS Genet, 10(1), e1004037.

Neverov, A. D., Kryazhimskiy, S., Plotkin, J. B., and Bazykin, G. A. (2015). Coordinated evolution of influenza a surface proteins. PLoS Genet, 11(8), e1005404.

Nguyen, L.-T., Schmidt, H. A., Von Haeseler, A., and Minh, B. Q. (2015). Iq-tree: a fast and effective stochastic algorithm for estimating maximum-likelihood phylogenies. Molecular biology and evolution, 32(1), 268–274.

Nik-Zainal, S., Van Loo, P., Wedge, D. C., Alexandrov, L. B., Greenman, C. D., Lau, K. W., Raine, K., Jones, D., Marshall, J., Ramakrishna, M., Shlien, A., Cooke, S. L., Hinton, J., Menzies, A., Stebbings, L. A., Leroy, C., Jia, M., Rance, R., Mudie, L. J., Gamble, S. J., Stephens, P. J., McLaren, S., Tarpey, P. S., Papaemmanuil, E., Davies, H. R., Varela, I., McBride, D. J., Bignell, G. R., Leung, K., Butler, A. P., Teague, J. W., Martin, S., Jönsson, G., Mariani, O., Boyault, S., Miron, P., Fatima, A., Langerød, A., Aparicio, S. A. J. R., Tutt, A., Sieuwerts, A. M., Borg, Å., Thomas, G., Salomon, A. V., Richardson, A. L., Børresen-Dale, A.-L., Futreal, P. A., Stratton, M. R., Campbell, P. J., and Breast Cancer Working Group of the International Cancer Genome Consortium (2012). The life history of 21 breast cancers. Cell, 149(5), 994–1007.

Posada, D. and Crandall, K. A. (2001). Intraspecific gene genealogies: trees grafting into networks. Trends in ecology & evolution, 16(1), 37–45.

Posada, D. and Crandall, K. A. (2002). The effect of recombination on the accuracy of phylogeny estimation. Journal of molecular evolution, 54(3), 396–402.

Powell, J. D., Abente, E. J., Chang, J., Souza, C. K., Rajao, D. S., Anderson, T. K., Zeller, M. A., Gauger, P. C., Lewis, N. S., and Vincent, A. L. (2020). Characterization of contemporary 2010.1 h3n2 swine influenza a viruses circulating in united states pigs. Virology, 553, 94–101.

Price, M. N., Dehal, P. S., and Arkin, A. P. (2010). Fasttree 2–approximately maximum-likelihood trees for large alignments. PloS one, 5(3), e9490.

Rajão, D. S., Gauger, P. C., Anderson, T. K., Lewis, N. S., Abente, E. J., Killian, M. L., Perez, D. R., Sutton, T. C., Zhang, J., and Vincent, A. L. (2015). Novel reassortant human-like H3N2 and H3N1 influenza A viruses detected in pigs are virulent and antigenically distinct from swine viruses endemic to the united states. Journal of Virology, 89(22), 11213–11222.

Rajão, D. S., Walia, R. R., Campbell, B., Gauger, P. C., Janas-Martindale, A., Killian, M. L., and Vincent, A. L. (2017). Reassortment between swine H3N2 and 2009 pandemic H1N1 in the United States resulted in influenza A viruses with diverse genetic constellations with variable virulence in pigs. Journal of virology, 91(4), e01763–16.

Rajão, D. S., Anderson, T. K., Kitikoon, P., Stratton, J., Lewis, N. S., and Vincent, A. L. (2018). Antigenic and genetic evolution of contemporary swine H1 influenza viruses in the United States. Virology, 518, 45–54.

Rasmussen, M. D. and Kellis, M. (2011). A bayesian approach for fast and accurate gene tree reconstruction. Molecular Biology and Evolution, 28(1), 273–290.

Robinson, D. and Foulds, L. (1981). Comparison of phylogenetic trees. Mathematical Biosciences, 53, 131–147.

Sagulenko, P., Puller, V., and Neher, R. A. (2018). Treetime: Maximum-likelihood phylodynamic analysis. Virus evolution, 4(1), vex042.

Scholtissek, C. (1990). Pigs as ‘mixing vessels’ for the creation of new pandemic influenza a viruses. Medical Principles and Practice, 2(2), 65–71.

Scholtissek, C., Bürger, H., Kistner, O., and Shortridge, K. (1985). The nucleoprotein as a possible major factor in determining host specificity of influenza h3n2 viruses. Virology, 147(2), 287–294.

Smith, G. J., Vijaykrishna, D., Bahl, J., Lycett, S. J., Worobey, M., Pybus, O. G., Ma, S. K., Cheung, C. L., Raghwani, J., Bhatt, S., et al. (2009). Origins and evolutionary genomics of the 2009 swine-origin H1N1 influenza A epidemic. Nature, 459(7250), 1122.

Solís-Lemus, C. and Ané, C. (2016). Inferring phylogenetic networks with maximum pseudolikelihood under incomplete lineage sorting. PLOS Genetics, 12(3), 1–21.

Steel, M. and Rodrigo, A. (2008). Maximum likelihood supertrees. Syst Biol, 57(2), 243–50.

Vaughan, T. G. (2017). Icytree: rapid browser-based visualization for phylogenetic trees and networks. Bioinformatics, 33(15), 2392–2394.

Venkatesh, D., Poen, M. J., Bestebroer, T. M., Scheuer, R. D., Vuong, O., Chkhaidze, M., Machablishvili, A., Mamuchadze, J., Ninua, L., Fedorova, N. B., et al. (2018). Avian influenza viruses in wild birds: virus evolution in a multihost ecosystem. Journal of virology, 92(15).

Wang, X., Minasov, G., and Shoichet, B. K. (2002). Evolution of an antibiotic resistance enzyme constrained by stability and activity trade-offs. Journal of molecular biology, 320(1), 85–95.

Weinreich, D. M., Delaney, N. F., DePristo, M. A., and Hartl, D. L. (2006). Darwinian evolution can follow only very few mutational paths to fitter proteins. science, 312(5770), 111–114.

Wen, D., Yu, Y., Hahn, M. W., and Nakhleh, L. (2016). Reticulate evolutionary history and extensive introgression in mosquito species revealed by phylogenetic network analysis. Molecular ecology, 25(11), 2361–2372.

Whidden, C., Beiko, R. G., and Zeh, N. (2013). Fixed-parameter algorithms for maximum agreement forests. SIAM Journal on Computing, 42(4), 1431–1466.

White, M. C. and Lowen, A. C. (2018). Implications of segment mismatch for influenza a virus evolution. The Journal of general virology, 99(1), 3.

White, M. C., Tao, H., Steel, J., and Lowen, A. C. (2019). H5n8 and h7n9 packaging signals constrain ha reassortment with a seasonal h3n2 influenza a virus. Proceedings of the National Academy of Sciences, 116(10), 4611–4618.

Willyard, A., Cronn, R., and Liston, A. (2009). Reticulate evolution and incomplete lineage sorting among the ponderosa pines. Molecular Phylogenetics and Evolution, 52(2), 498–511.

Woolley, S. M., Posada, D., and Crandall, K. A. (2008). A comparison of phylogenetic network methods using computer simulation. PLoS One, 3(4), e1913.

Yu, Y. and Nakhleh, L. (2015). A maximum pseudo-likelihood approach for phylogenetic networks. BMC genomics, 16(10), 1–10.

Yu, Y., Degnan, J. H., and Nakhleh, L. (2012). The probability of a gene tree topology within a phylogenetic network with applications to hybridization detection. PLoS genetics, 8(4), e1002660.

Yu, Y., Barnett, R. M., and Nakhleh, L. (2013). Parsimonious inference of hybridization in the presence of incomplete lineage sorting. Systematic Biology, 62(5), 738–751.

Zeller, M. A., Anderson, T. K., Walia, R. W., Vincent, A. L., and Gauger, P. C. (2018). ISU FLUture: a veterinary diagnostic laboratory web-based platform to monitor the temporal genetic patterns of Influenza A virus in swine. BMC Bioinformatics, 19(1), 397.

Zeller, M. A., Chang, J., Vincent, A. L., Gauger, P. C., and Anderson, T. K. (2020). Coordinated evolution between n2 neuraminidase and h1 and h3 hemagglutinin genes increased influenza a virus genetic diversity in swine. bioRxiv.

Zhang, Y., Aevermann, B., Anderson, T., Burke, D., Dauphin, G., Gu, Z., He, S., Kumar, S., Larsen, C., Lee, A., et al. (2017). Influenza research database: An integrated bioinformatics resource for influenza virus research. Nucleic acids research, 45(D1), D466.

